# T cell polarization and NFAT translocation are stiffness-dependent and are differentially regulated by Piezo1 and Orai1

**DOI:** 10.1101/2024.03.06.583253

**Authors:** Renping Zhao, Jingnan Zhang, Eva C. Schwarz, Aránzazu del Campo, Markus Hoth, Bin Qu

## Abstract

Effective T cell responses against tumor cells require diverse effector functions including polarization towards tumor cells to form immunological synapses and nuclear factor of activated T-cells (NFAT)-dependent gene transcription. While the role of tumor cell softening has been associated with malignancy, stemness, and metastasis, potentially contributing to immune evasion, its impact on cellular processes in T cells is not well understood. Here, we show that both T cell polarization and NFAT nuclear translocation are modulated by target stiffness in a Ca2+ dependent manner. Using both anti-CD3 antibody-functionalized substrates with varying stiffness as surrogates for target cells or softened tumor cells, we found that both, reorientation of microtubule organizing center (MTOC) towards the tumor cells, a hallmark for T cell polarization, and NFAT translocation were impaired on softer hydrogels or following contact with softer cancer cells. The amplitudes of intracellular Ca2+ signals were dependent on stiffness, and removal of extracellular Ca2+ inhibited stiffness-dependent T cell responsiveness. While stiffness-dependent Ca2+ signaling was crucial for both, T cell polarization and NFAT translocation, Ca2+ influx through Piezo1, a mechanosensitive ion channel, mediated stiffness-dependent MTOC reorientation but not NFAT translocation. In contrast, Ca2+ influx through store-operated Orai channels mediated NFAT translocation but not MTOC reorientation. Our results demonstrate that tumor cell stiffness directly influences T cell functionality through distinct Ca2+ influx pathways, revealing cell softening as an essential mechanism employed by malignant cells to evade immune surveillance.

## Introduction

One significant challenge that immune cells face within the solid tumor microenvironment is the phenomenon of cancer cell softening, a characteristic linked to increased malignancy, metastasis, and stemness. Compelling evidence at the single-cell level, derived from both optical stretching methods and atomic force microscopy, highlights that carcinoma cells, including those from breast, cervix, and ovarian cancers, exhibit greater deformability with lower Young’s moduli in comparison to their non-malignant counterparts (1-4). Among cancerous cells, those displaying softer mechanical properties exhibit increased invasiveness, higher potentials to establish metastatic clones, and an up-regulation of gene profiles associated with stemness (2, 5, 6). Moreover, when malignant cancer cells navigate confined channels, they can undergo further softening (6). However, the extent to which cancer cell softening facilitates the evasion of immune surveillance by malignant cells remains poorly understood.

As a crucial component of immune responses against cancer, T cells must promptly and robustly respond to stimuli, with activation needing to be precisely tuned, either propagated or terminated, to effectively eliminate cancer cells without compromising healthy cells or tissues. Upon identifying cognate target cells, the cytoskeleton of T cells undergoes polarization towards these targets, ensuring the specific delivery of vesicles containing cytokines or cytotoxic proteins (7, 8). A fundamental rearrangement of the cytoskeleton involves the rapid translocation of the microtubule-organizing center (MTOC) towards the target cells, acting as a hallmark for the formation of the immunological synapse (IS) and playing a pivotal role in subsequent effector functions (9-11). In T cells, the failure of MTOC reorientation towards the IS is associated with the lack of enrichment of lytic granules, significantly impaired lytic capability (12, 13), and reduced IL-2 production (14). For further T cell commitment, gene regulation by nuclear factor of activated T cells (NFAT) plays a central role in T cell fate determination, either sustaining T cells in an activated state or reprogramming T cells into an unresponsive state (15, 16). NFAT activity depends on its translocation from the cytosol into the nucleus following TCR activation (15-17).

The elevation of intracellular calcium (Ca^2+^) levels ([Ca^2+^]_int_) is a hallmark of T cell activation, and the resulting [Ca^2+^]_int_ increase triggered by target recognition plays a pivotal role in both NFAT nuclear translocation and MTOC polarization. In the resting state, NFAT resides in the cytosol in its phosphorylated form; upon activation, Ca^2+^-dependent calcineurin dephosphorylates NFAT proteins, enabling their translocation to the nucleus (18-20). Deficiencies in store-operated Ca^2+^ entry (SOCE) in T cells impede NFAT activation, hindering nuclear translocation following IS formation (21, 22). Although the role of Ca^2+^ in MTOC polarization is not entirely elucidated, the absence of extracellular Ca^2+^ has been observed to diminish MTOC translocation in response to target cells (23, 24). Interestingly, chelation of both extracellular and intracellular Ca^2+^ significantly impairs MTOC translocation (10) but removal of intracellular Ca^2+^ with BAPTA does not affect MTOC repositioning (13).

Store-operated Ca^2+^ entry (SOCE) is the primary Ca^2+^ influx mechanism in T cells (25, 26). Upon the recognition of target cells, TCR activation initiates the production of inositol trisphosphate (IP_3_), which binds to its receptor IP_3_R at the endoplasmic reticulum (ER), prompting the efflux of Ca^2+^ from the ER into the cytosol (27). Ca^2+^ depletion in the ER is sensed by stromal interaction molecules (STIM) and the clustering of STIM in the ER membrane activates calcium release-activated calcium (CRAC) channels called Orai, allowing the entry of extracellular Ca^2+^ (28, 29). Defects in Orai proteins in patients are associated with severe immunodeficiency due to the failure of T cell activation (30-32).

Ca^2+^ entry can also be mediated by other ion channels, including Piezo1. Piezo1 is a trimeric mechanosensitive cation channel and is permeable to Ca^2+^ (33, 34). In T cells, the activation of Piezo1 induces Ca^2+^ influx, and the downregulation of Piezo1 hampers the activation and proliferation of T cells (35). *In vivo*, Piezo1 deficiency promotes the expansion of regulatory T cells, thereby attenuating the severity of experimental autoimmune encephalomyelitis, a mouse model for multiple sclerosis (36). Piezo1 is also implicated in the regulation of integrin-dependent chemotaxis (37) and the shear stress-enhanced activation of T cells (38).

In this study, we show that decreased substrate stiffness or softer tumor cells impair the responsiveness of T cells, as evident from Ca^2+^ influx, NFAT nuclear translocation, and MTOC reorientation towards the IS. Both Orai1 and Piezo1 contribute to the IS formation-triggered Ca^2+^ entry in a stiffness-dependent manner. Piezo1 plays a crucial role in the rapid increasing phase, while Orai1 is more important for the sustainability of elevated intracellular Ca^2+^ concentrations. Orai1-mediated Ca^2+^ entry is required for NFAT nuclear translocation, as expected from the literature (21, 22) but, unexpectedly, not for MTOC reorientation. On the opposite, Piezo1-mediated fast Ca^2+^ increases primarily influence MTOC reorientation, with no discernible impact on NFAT nuclear translocation. These findings assign specific functions to Ca^2+^ influx by Orai and Piezo1, unveiling a hitherto unreported intrinsic mechanism that coordinates the rapid and long-term cellular responses to stiffness cues.

## Results

Several key processes collectively optimize and modulate efficient T cell responses against pathogens or cancer. Among those, MTOC reorientation to the immunological synapse (IS) is a pivotal event during T cell polarization, and NFAT nuclear translocation is essential to initiate gene transcription programs. Given that T cells are exposed to varying levels of stiffness in their microenvironment, we analyzed how different stiffness levels of 2, 16, and 64 kPa, covering the physiological range (39), influence T cell responses. We used the human T cell line Jurkat as a model system and settled them on functional hydrogels of different stiffness (Fig. 1a), which were coated with anti-CD3 antibodies to activate T cell receptors (TCRs), the main initiators of IS formation and T cell signaling. T cells were fixed at different time points, and we analyzed MTOC localization or NFAT translocation. In Fig. 1b and Supplementary Fig. 1, it is evident that MTOC, stained with anti-tubulin antibody, was consistently closer to the IS at all time points at medium (16 kPa) and stiff (64 kPa) substrates compared to soft ones (2 kPa). Similarly, NFAT translocation to the nucleus stained with Hoechst was fastest at 64 kPa, as observed in the respective fluorescence images and the corresponding quantification (Fig. 1c, d). Similar stiffness-dependent responsiveness was observed in primary human CD4^+^ T cells (Fig. 1e, f). Given the amounts of anti-CD3 antibodies on the different surfaces were very similar for 2, 16, and 64 kPa (Supplementary Fig. 2), the stiffness of the substrates is the sole differing parameter under these experimental conditions. In addition, substrate-functionalized CD28 antibodies did not induce MTOC reorientation or NFAT nuclear translocation (Supplementary Fig. 3). Our results thus suggest that substrate stiffness modulates both, MTOC reorientation and NFAT nuclear translocation in a CD3 activation dependent manner.

**Figure 1.**
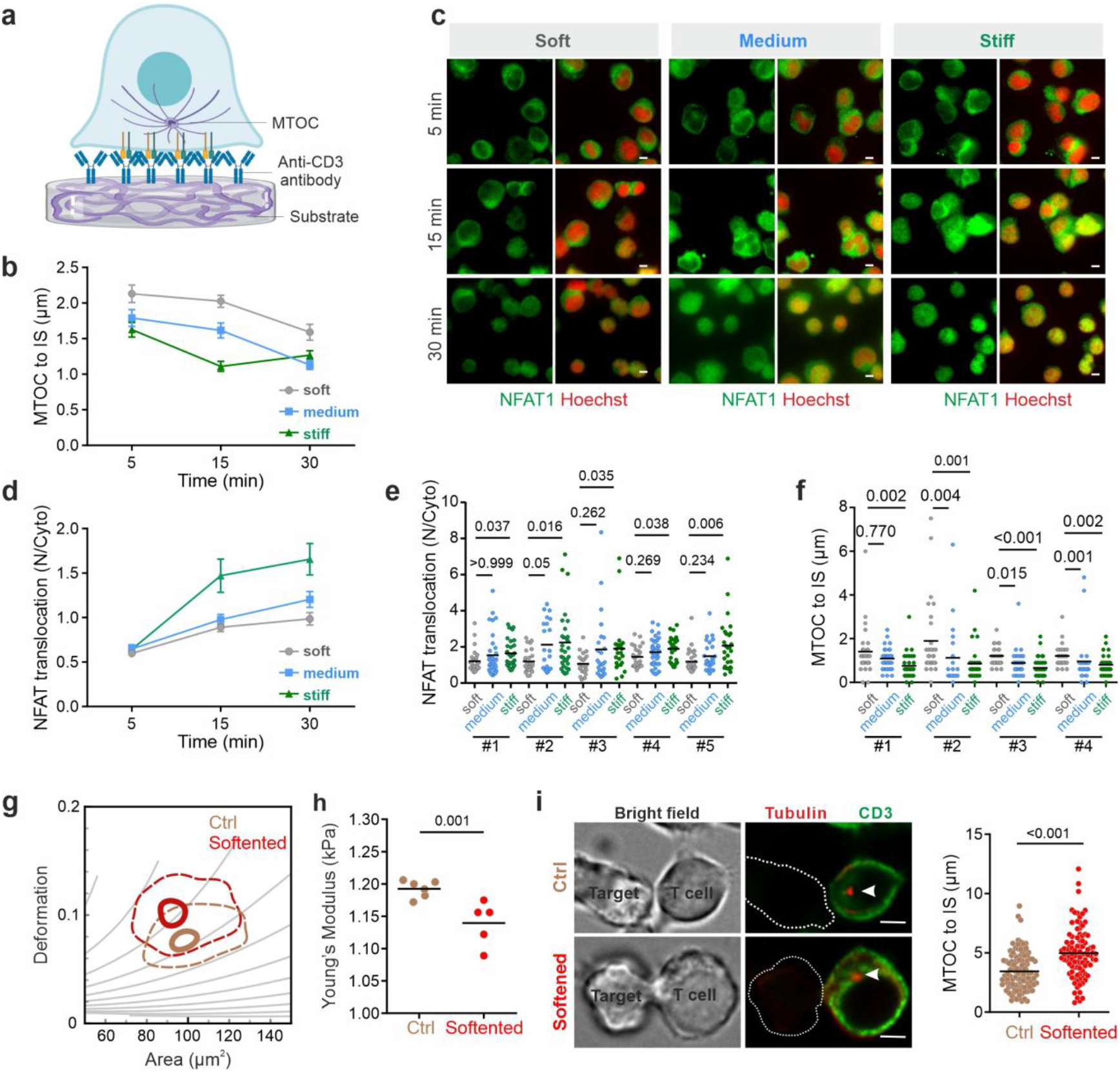
The impact of target cell stiffness on MTOC reorientation and NFAT nuclear translocation. (**a**) Schematic of anti-CD3 antibody-functionalized hydrogel with T cell contact. (**b-d**) Stiffness influences MTOC reorientation and NFAT translocation in Jurkat T cells on substrates. Jurkat T cells were settled on functionalized substrates for varying time periods at 37°C before fixation. The results are from 3 independent experiments. (**b**) MTOC was stained with the anti-Tubulin antibody and the position of the IS was identified with Phalloidin-labeled actin. (**c, d**) NFAT and the nucleus were labeled with anti-NFAT antibody and Hoechst 33342, respectively. (**e-f**) Stiffness influences MTOC reorientation and NFAT translocation in T-Activator CD3/CD28 antibody beads stimulated human primary CD4^+^ T cells on substrates. Human primary CD4^+^ T cells were settled on functionalized substrates for 15 min at 37°C before fixation. (**e**) NFAT and the nucleus were labeled with anti-NFAT antibody and Hoechst 33342, respectively. The results are from 5 donors. (**f**) MTOC was stained with the anti-Tubulin antibody and the position of the IS was identified with Phalloidin-labeled actin. The results are from 4 donors. (**g-h**) EGTA-AM treatment softens cancer cells. NALM-6 cells were pre-treated with 500 µM of EGTA-AM for 30 min at 37°C. Cell stiffness was measured with RT-DC. One representative experiment is shown in **g** and the quantification is shown in **h**, in which each dot represents the mean value of one independent experiment. The linear mixed model was conducted for statistical significance. (**i**) Softening of cancer cells hinders MTOC translocation. NALM-6 cells were pre-treated with 500 µM EGTA-AM for 30 min at 37°C. Jurkat T cells were conjugated with target cells at 37°C for 15 min before fixation. The MTOC was stained with anti-tubulin antibody and indicated with white arrowheads. 96-well CytoSoft plates were used for soft (2 kPa), medium (16 kPa), and stiff (64 kPa) substrates. Student’s t-test (with two tails) was conducted for **i**. The data are shown as mean±SEM in **b** and **d**. Dunn’s multiple comparisons test (with two tails) was conducted for **e**-**f**. Scale bars are 5 µm.

We further explored the impact of tumor cell softening on T cell responses. We softened NALM-6 tumor cells by chelating intracellular Ca^2+^ with EGTA-AM. The stiffness of NALM-6 cells was assessed using real-time deformability cytometry (RT-DC), a high throughput technique that quantifies cell deformation along with cell size as cells pass through narrow channels (40). In comparison to control cells, EGTA-AM-loaded NALM-6 cells exhibited slightly enhanced deformation (Fig. 1g) and correspondingly moderately reduced Young’s moduli (Fig. 1h). We conjugated softened NALM-6 tumor cells with T cells and analyzed MTOC reorientation in T cells. We found that the moderate reduction in the stiffness of EGTA-loaded target cells significantly increased the distance between the MTOC and the IS in the conjugated T cells (Fig. 1i), indicating impaired MTOC translocation towards the IS. Thus, we conclude that target cell stiffness plays a pivotal role in regulating T cell functionality triggered by TCR activation.

Given the crucial role of Ca^2+^ influx in NFAT translocation (19-22) and its modulation by stiffness as shown above, we hypothesized that stiffness-dependent changes in Ca^2+^ influx may also underlie the modulation of MTOC reorientation. For assessing intracellular Ca^2+^ levels ([Ca^2+^]_int_), we utilized functionalized polyacrylamide (PA) hydrogels prepared on coverslips to allow Ca^2+^ imaging. Instead of 2, 16 and 64 kPa, we opted for 2, 12 and 50 kPa, because these conditions have been optimized previously for comparable anti-CD3 antibody coating efficiency for this setting (41). The successful formation of a functional IS between T cells and the antibody-functionalized hydrogels was confirmed, as evident by F-actin localization, CD3 localization, and ZAP-70 phosphorylation (Supplementary Fig. 4). Regarding T cell responsiveness, we noted that stiffer hydrogels correlated with higher proportions of T cells displaying a responsive behavior, characterized by an elevation in [Ca^2+^]_int_ (Fig. 2a). In all T cells, [Ca^2+^]_int_ was substantially higher on stiffer hydrogels compared to those of medium and soft rigidity, as illustrated by three representative examples (Fig. 2b), and the average of all analyzed cells (Fig. 2c). This stiffness-dependent changes in Ca^2+^ responses are also reflected by rate of [Ca^2+^]_int_ increase (Fig. 2c-d), the peak value of [Ca^2+^]_int_ (Fig. 2e), and the plateau (Fig. 2f). For medium and high stiffness levels, [Ca^2+^]_int_ remained elevated throughout the measurement (Fig. 2c). While control experiments with CD28 antibody-coated hydrogels induced T cell spreading (Supplementary Fig. 5) to similar levels as seen with CD3 antibody-functionalized hydrogels, no [Ca^2+^]_int_ increase on any of the CD28 antibody-functionalized substrates (Supplementary Fig. 6). This confirms that Ca^2+^ signals were triggered by CD3 engagement and TCR activation, rather than simply by T cell adhesion to the surface. In addition, we found that inhibiting Ca^2+^ influx by chelating extracellular Ca^2+^ significantly compromised TCR activation-induced MTOC reorientation (Fig. 2g). Collectively, our data suggest that [Ca^2+^]_int_ increases by Ca^2+^ influx plays a critical role in regulating T cell functionality in response to stiffness cues.

**Figure 2.**
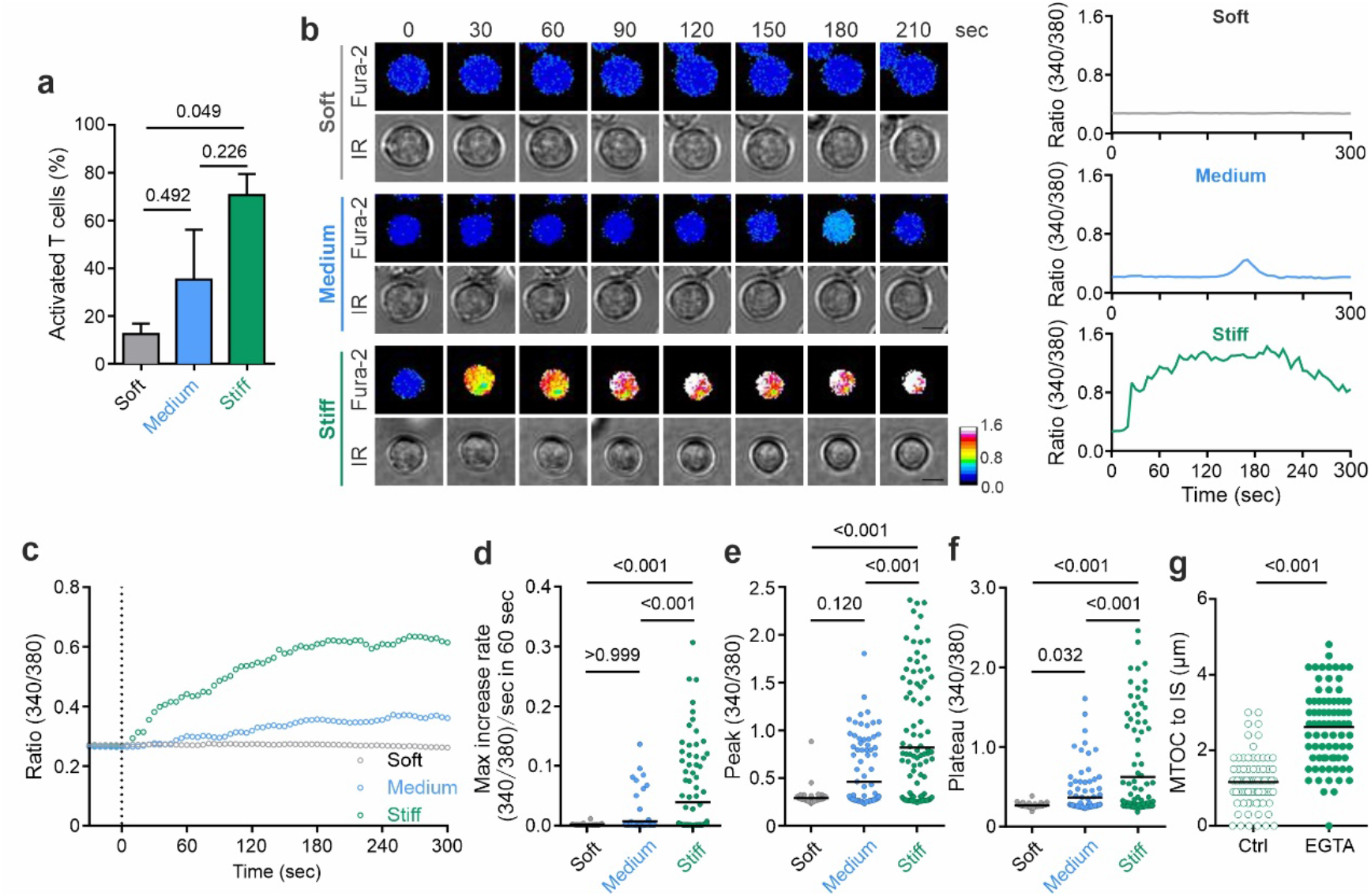
TCR activation triggered Ca^2+^ rise is modulated by substrate stiffness and is required for MTOC reorientation. (**a-f**) Jurkat T cells were loaded with Fura-2 and settled on anti-CD3 antibody functionalized hydrogels. Live-cell imaging with 340 nm, 380 nm, and infrared (IR) was acquired every 5 sec at RT. The corresponding Ca^2+^ signals are presented in **a**; IR: Infrared. The T cells showing elevated [Ca^2+^]_int_ after gel contact are shown in **b**. Scale bars are 5 µm. Mean values of all cells are shown in **c**, in which time 0 is defined as the time point when T cells settle down on the surface. Ca^2+^ max increase rates (**d**), peak (**e**), and plateau (**f**) were shown in the figure. (**g**) Extracellular Ca^2+^ depletion hinders MTOC reorientation. Jurkat T cells were settled on anti-CD3 antibody functionalized stiff substrates (64 kPa) in the presence or absence of EGTA (0.5 mM) for 15 min before fixation. MTOC was stained with the anti-tubulin antibody and the position of the IS was identified with phalloidin-labeled actin. Results are from three independent experiments. A Krushal-Wallis test with Dunn’s multiple comparisons was conducted for **d, f**. A one-way ANOVA test with Tukey’s multiple comparisons and the Mann-Whitney test was conducted for **a, e**, and **g**, respectively. Functionalized hydrogels with stiffness levels of 2 kPa (soft), 12 kPa (medium), and 50 kPa (stiff) were used in a-f. 96-well CytoSoft plates with a stiffness level of 64 kPa (stiff) were used in g. All the results are from at least three independent experiments.

To identify the underlying Ca^2+^ channels, we first focused on Orai1 as a potential candidate. Orai1 is the predominantly expressed Ca^2+^ release-activated Ca^2+^ (CRAC) channel in T cells and mediates the major proportion of Ca^2+^ influx. It is well-established that Orai1-dependent Ca^2+^ influx governs NFAT nuclear translocation and activity in T cells (42-44). we utilized Jurkat T cells to investigate the regulatory role of Orai1 in stiffness-dependent T cell responses, in which the Orai1 expression levels are comparable to primary human CD4^+^ T cells (Supplementary Fig. 7). To inhibit Orai1 function, we employed BTP-2 (2-aminoethyl diphenylborinate), a dose-dependent inhibitor primarily targeting CRAC/Orai1 channels (45). We verified that at a very high concentration (10 µM), BTP2 completely eliminated SOCE, without affecting [Ca^2+^]_int_ levels of ER store depletion (Supplementary Fig. 8) as reported previously (45, 46). Interestingly, while the inhibition of Orai1 channels by BTP2 completely abolished NFAT nuclear translocation for all three substrate stiffness levels (Fig. 3a), it had no impact on MTOC reorientation or its modulation by stiffness (Fig. 3b). To further confirm the role of Orai1, we employed siRNA to down-regulate Orai1 expression (Supplementary Fig. 9) and indeed observed that down-regulation of Orai1 in Jurkat T cells inhibited stiffness-dependent NFAT nuclear translocation and showed no effect on MTOC reorientation (Fig. 3c-d). Regarding CD3-induced Ca^2+^ signals, surprisingly, we found that they were not completely inhibited by BTP2 on stiff substrates (50 kPa). Quantification revealed that BTP2 application under these conditions did not alter [Ca2+]_int_ increase rate (Fig. 3f) and peak values of [Ca2+]_int_ (Fig. 3g) but reduced [Ca^2+^]_int_ over longer times (Fig. 3h). A similar effect of BTP2 was also found for medium stiffness (12 kPa, Supplementary Fig. 10). Interestingly, expression level of Orai 1 is considerably lower in unstimulated primary human CD4^+^ T cells compared to their stimulated counterparts (Supplementary Fig. 11a), and NFAT nuclear translocation did not exhibit a stiffness-dependent manner (Supplementary Fig. 11b), again indicating that Orai1 expression levels are crucial for the stiffness dependence of NFAT translocation. Our findings suggest that Orai1 plays a critical role in activated T cells for stiffness-regulated NFAT nuclear translocation, however, another stiffness-regulated Ca^2+^ influx pathway must mediate MTOC reorientation.

**Figure 3.**
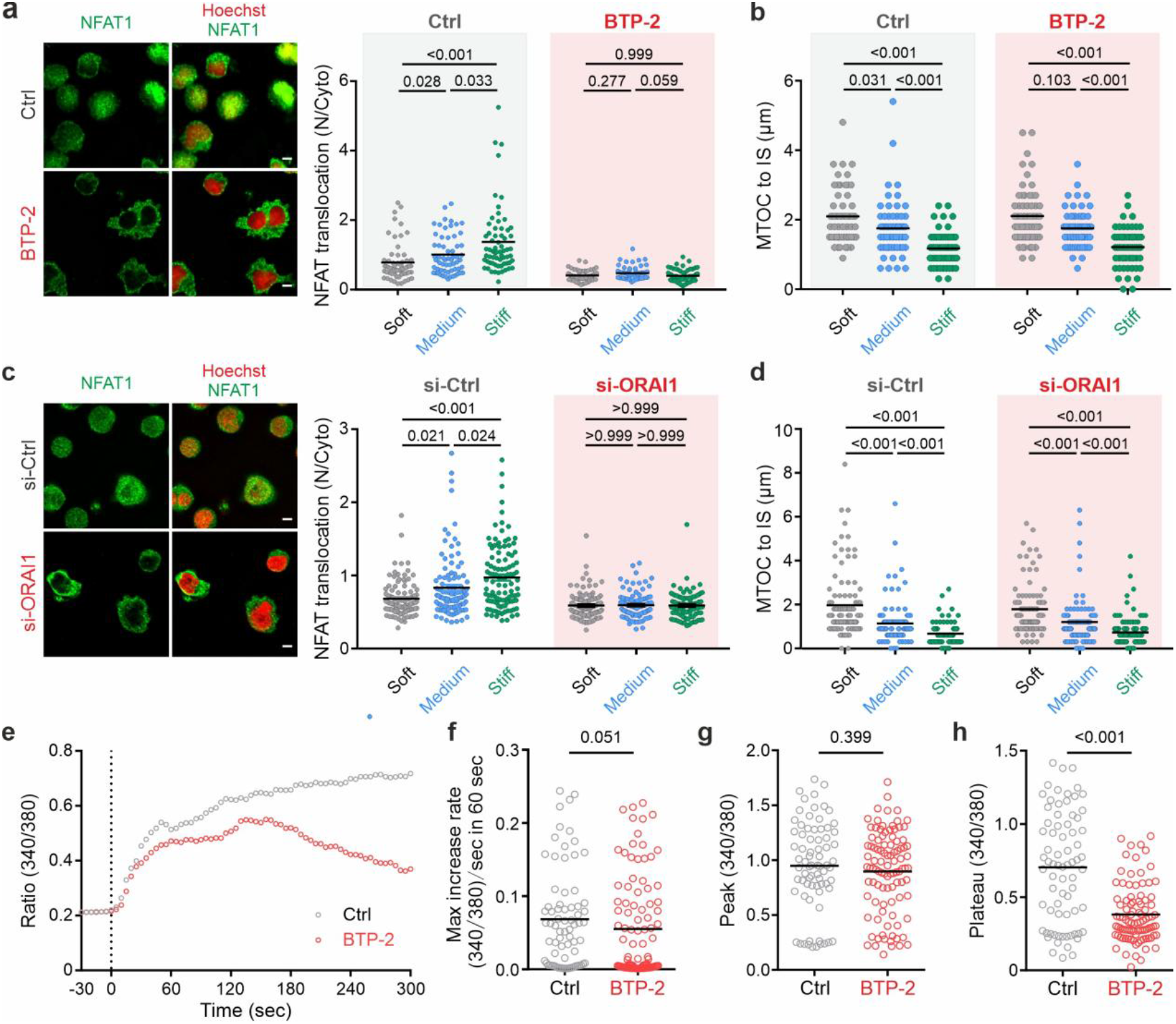
CRAC channels are crucial for stiffness-dependent NFAT nuclear translocation and the maintenance of elevated [Ca^2+^]_int_. (**a**-**d**) Orai1 governs stiffness dependent NFAT translocation, but not involved in MTOC reorientation. (**a**-**b**) Jurkat T cells were pre-treated with BTP-2 (10 µM) at 37°C for 30 min. (**a**) NFAT and the nucleus were stained with anti-NFAT antibody and Hoechst 33342, respectively. Exemplary cells on stiff substrates are shown. (**b**) MTOC was stained with the anti-tubulin antibody and the position of the IS was identified with Phalloidin-labeled actin. (**c**-**d**) Orai1 was knocked down in Jurkat T cell with siRNA. T cells were plated on anti-CD3 antibody-functionalized surface at 37°C for 15 min before fixation. (**c**) NFAT and the nucleus were stained with anti-NFAT antibody and Hoechst 33342, respectively. Exemplary cells on stiff substrates are shown. (**d**) MTOC was stained with the anti-tubulin antibody and the position of the IS was identified with Phalloidin-labeled actin. (**e**-**h**) Impact of Orai1 channels on TCR activation-triggered Ca^2+^ rise. Jurkat T cells were settled on stiff substrates (50 kPa hydrogel). The mean values of all cells for each condition are shown in **e**, in which time 0 is defined as the time point when T cells settle down on the surface. Ca^2+^ max increase rates (**f**), peak (**g**), and plateau (**h**) were quantified. Scale bars are 5 µm. All results are from three independent experiments. A Krushal-Wallis test with Dunn’s multiple comparisons test was conducted for **a, b**. A Mann-Whitney test was conducted for **e-g**. 96-well CytoSoft plates were used for soft (2 kPa), medium (16 kPa), and stiff (64 kPa) substrates. All the results are from at least three independent experiments.

Mechanosensitive ion channels are crucial for adapting T cell behaviour to mechanical cues and Piezo1 is Ca^2+^ permeable and also important for T cell functions (39, 47). Using flow cytometry, we identified that Piezo1 was expressed at a comparably high level in both Jurkat and primary human CD4^+^ T cells (Fig. 4a). To investigate whether Piezo1 regulates Ca^2+^ entry in T cells, we downregulated Piezo1 using siRNA (Fig. 4b). Its downregulation resulted in reduced Ca^2+^ influx, as shown in [Ca^2+^]_int_ increase rates (Fig. 4c), but not in peak levels (Fig. 4d), and plateau levels (Fig. 4e). In comparison, when T cell was activated with medium substrates (12 kPa), the difference between Piezo1 down-regulated and control Jurkat T cells was not statistically significant in terms of CD3-activation induced [Ca^2+]^_int_ levels (Supplementary Fig. 12). Interestingly, compared to their stimulated counterparts, unstimulated primary human CD4^+^ T cells expressed considerably low levels of Piezo1 (Supplementary Fig. 13a), and their MTOC reorientation did not exhibit a stiffness-dependent manner (Supplementary Fig. 13b). These findings show that Piezo1 is involved in initiation of Ca^2+^ influx upon TCR activation in activated T cells and is mainly contributing to the fast-rising phase, especially when T cells are activated on stiffer substrates.

**Figure 4.**
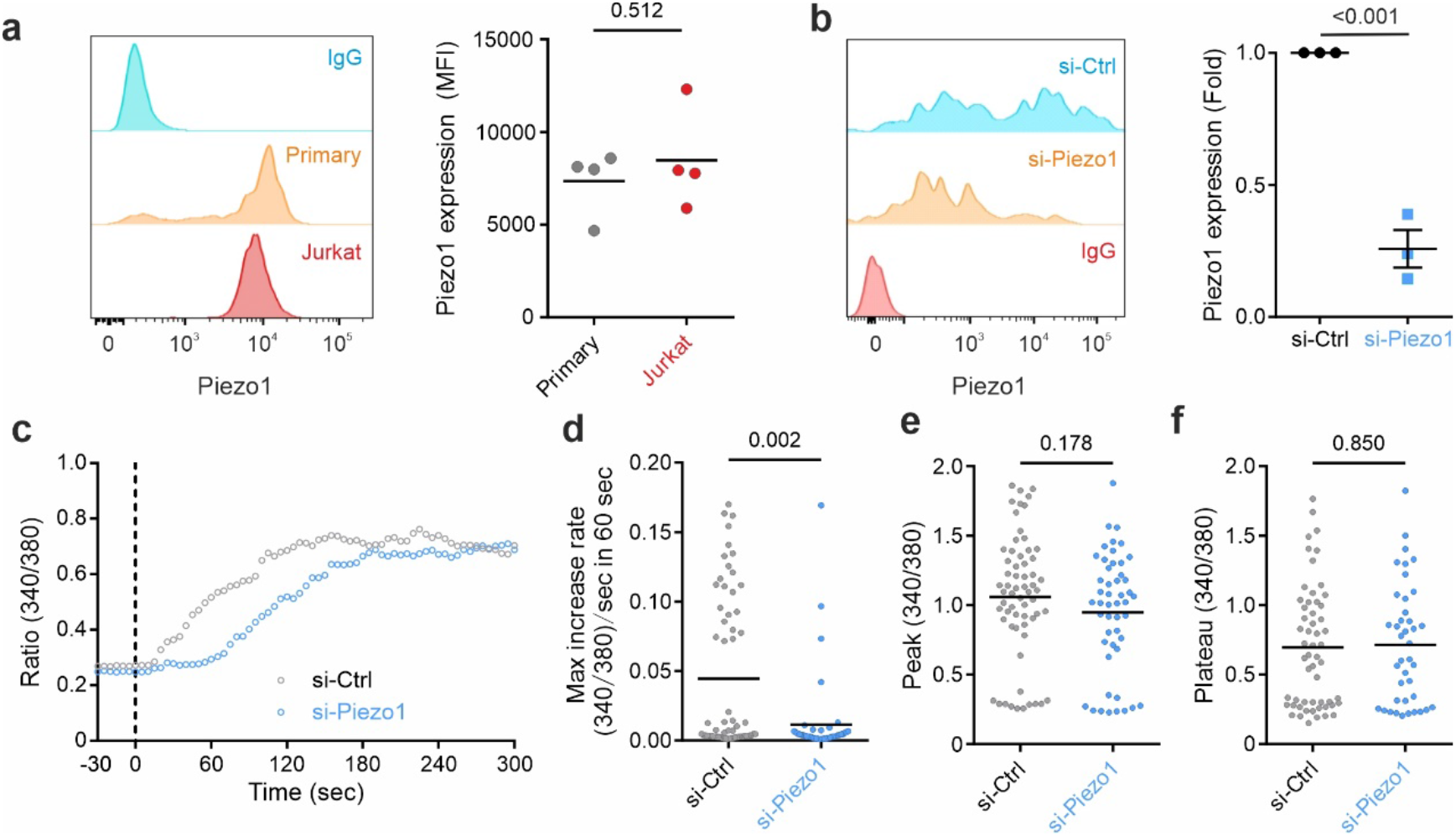
Piezo1 regulates Ca^2+^ increase rates. (**a**) Expression of Piezo1 in T-Activator CD3/CD28 antibody beads stimulated human primary CD4^+^ T cells and Jurkat T cells. Results are from four independent experiments. (**b**) Down-regulation of Piezo1 by siRNA. Jurkat T cells were transfected with control siRNA (si-Ctrl) or siRNA targeting Piezo1 (si-Pizeo1). Transfected cells were fixed 24 hours after transfection and stained with Piezo1 antibody. The expression of Piezo1 was measured with flow cytometry. IgG staining was used for setting the gate. One representative experiment is shown in the left panel. Quantification of fold change in mean fluorescence intensity is shown in the right panel. Results are from three independent experiments. (**c-f**) Impact of Piezo1 on TCR activation-triggered Ca^2+^ rise. Jurkat T cells transfected with siRNA were loaded with Fura-2-AM and then settled on anti-CD3 antibody functionalized stiff hydrogels (50 kPa). Averaged curves of cells for each condition are shown in **c**, in which time 0 is defined as the time point when T cells settle down on the surface. Ca^2+^ max increase rate (**d**), peak (**e**), and plateau (**f**) were quantified for individual cells. A student’s t-test was conducted for **a, b**. A Mann-Whitney test was conducted for **d-f**. All results are from three independent experiments in **c**-**f**.

Next, we examined the role of Piezo1 in T cell polarization. In clear contrast to the inhibition of Orai1 channels, inhibition of Piezo1, either by siRNA (Fig. 5a) or its inhibitor GsMTx4 (Fig. 5b), abolished the dependence of MTOC reorientation on stiffness. Similarly, in primary human CD4^+^ T cells, GsMTx4 treatment substantially impaired MTOC repositioning towards IS (Fig. 5c). Consistently, for T cells conjugated with superantigen SEA/SEB-pulsed target cells, perturbation of Piezo1, either by siRNA or GsMTx4, also impaired MTOC reorientation induced by target cell recognition as evident from the increase between MTOC and IS (Fig. 5d-e). In addition, activation of Piezo1 by its specific agonist Yoda-1 increased [Ca^2+^]_int_ (Fig. 5f), which significantly increased MTOC reorientation and NFAT nuclear translocation on soft substrates (Fig. 5g-h). Results from live-cell imaging show that inhibition of Piezo1 by GsMTx-4 delayed the initiation of MTOC translocation towards the IS and increased the minimal distance between MTOC and IS (Fig. 5i, j). In contrast, when CRAC/Orai1 channels were inhibited by BTP-2, MTOC reorientation was similar compared to control (Fig. 5i, j). Notably, NFAT translocation to the nucleus, was unaffected by Piezo1 (Fig. 5k). These findings suggest that Piezo1-mediated mechanosensing plays a critical role in MTOC reorientation and concomitant T cell polarization without interfering with NFAT nuclear translocation. Collectively, we conclude that Piezo1 governs dynamics of MTOC translocation in response to stiffness.

**Figure 5.**
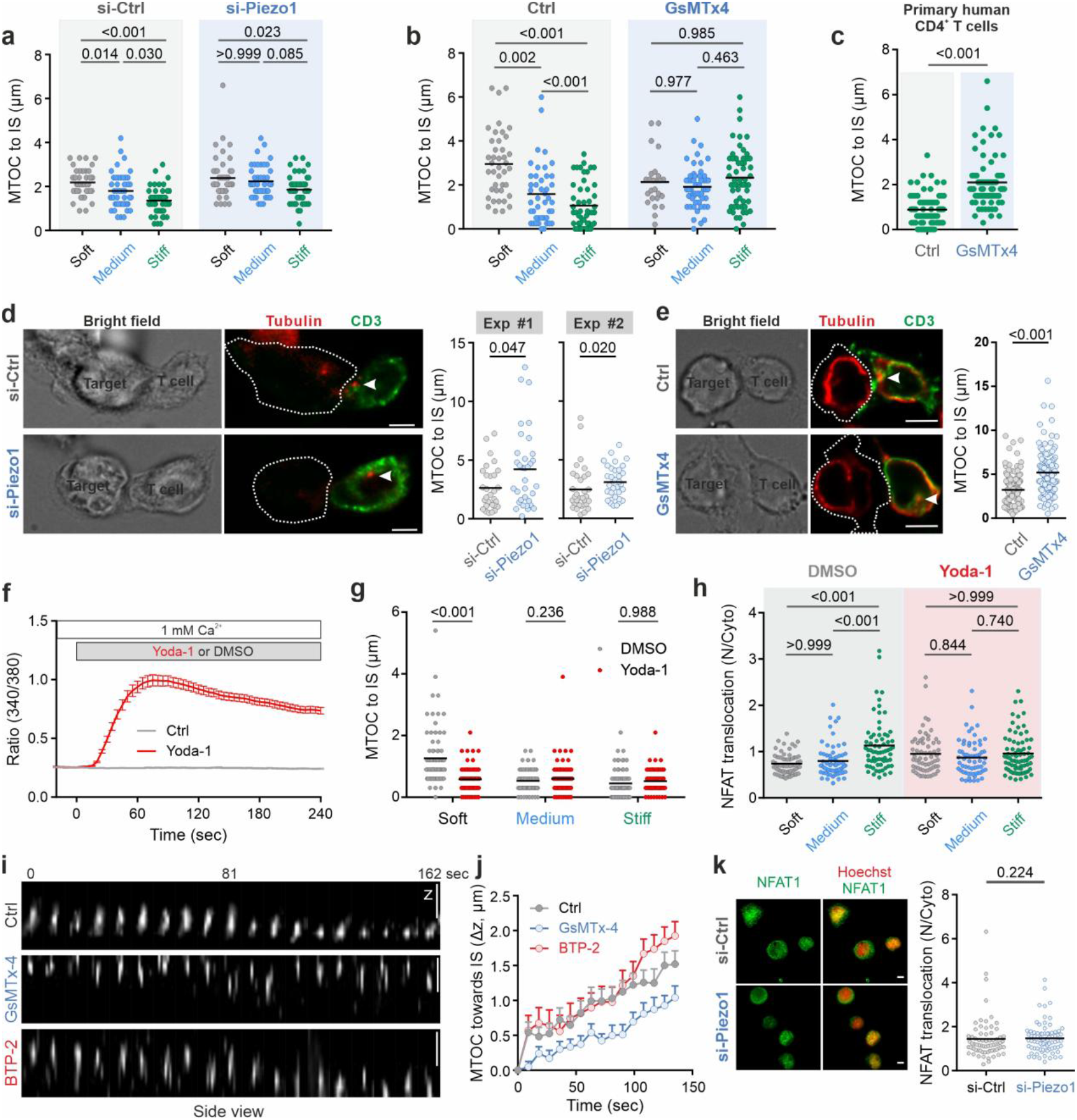
Piezo1 plays a pivotal role for MTOC reorientation but not for NFAT translocation. (**a-e**) Perturbation of Piezo1 hinders stiffness-dependent MTOC reorientation. Jurkat T cells (**a, b, d, e**) or primary human CD4^+^ T cells (**c**) were used. T cells were either transfected with siRNA (**a, d**) or treated with GsMTx-4 (1 µM) (**b, c, e**). In **a-c**, T cells were settled on anti-CD3-antibody functionalized substrates. In **d**-**e**, T cells were conjugated with SEA/SEB pulsed target cells, either NALM-6 cells (**d**) or Raji cells (**e**), at 37°C for 15 min before fixation. MTOC was stained with the anti-tubulin antibody. The position of the IS was identified with phalloidin-labeled actin. White arrowheads in **d** and **e** highlight the position of MTOC (**f**) Yoda-1 treatment induces Ca^2+^ influx in T cells. Jurkat T cells loaded with Fura-2-AM were settled on poly-L-ornithine-coated coverslips in 1 mM Ca^2+^ ringer solution. Yoda-1 (15 µM) was perfused at the time 0. (**g**-**h**) Activation of Piezo1 sensitizes MTOC reorientation on soft substrates (**g**) but does not impact NFAT nuclear translocation (**h**). Jurkat T cells were settled on anti-CD3-antibody functionalized substrates in the presence of Yoda-1 (15 µM) at 37°C for 15 min before fixation. MTOC was stained with anti-Tubulin antibodies, and T cell contour was indicated with Phalloidin-labeled actin (**g**). NFAT and the nucleus were stained with anti-NFAT antibody and Hoechst 33342, respectively (**h**). (**i**-**j**) Piezo1 but not CRAC/Orai1 governs MTOC reorientation. Jurkat T cells were pre-treated with GsMTx-4 (1 µM), BTP-2 (10 µM) or vehicle control (Ctrl) at 37°C for 30 min. MTOC was stained by SPY650-tublin and visualized every 9 sec for 30 min at RT on anti-CD3-antibody-functionalized stiff substrates (64 kPa). (**k**) Piezo1 has no impact on NFAT nuclear translocation. Jurkat T cells transfected with siRNA were settled on anti-CD3-antibody-functionalized stiff substrates (64 kPa) at 37°C for 15 min before fixation. NFAT was labeled using anti-NFAT antibodies, and the nucleus was labeled with Hoechst 33342. A Krushal-Wallis test with Dunn’s multiple comparisons was conducted for **a**. A one-way ANOVA with Tukey’s multiple comparisons was conducted for **b**. A Mann-Whitney test was conducted for **c, d, e**, and **j**. In **a, g, h**, 96-well CytoSoft plates with stiff levels of 2 kPa (soft), 16 kPa (medium), and 64 kPa (stiff) were used. In **b**, functionalized hydrogels with stiffness levels of 2 kPa (soft), 12 kPa (medium), and 50 kPa (stiff) were used. In **c** and **h-j**, 96-well CytoSoft plates with 64 kPa (stiff) were used. Scale bars are 5 µm. In all scatter dot plots, one dot indicates one cell. In **i**, the data are shown as mean±SEM (Ctrl: 45 cells, GsmTx-4: 45 cells, BTP-2: 39 cells). Results in (**d, f**) were from two independent experiments. All other results were from three independent experiments or three donors. Scale bars are 5 µm.

In summary, our findings establish Piezo1 as a mechanosensor in T cell, governing MTOC reorientation and concurrent polarization. However, its Ca^2+^ signals do not contribute to NFAT translocation. In contrast, beyond its known role in guiding NFAT translocation (21, 22), Orai1 serves as a key regulator for stiffness-dependent modulation of NFAT translocation but is not involved in MTOC reorientation.

## Discussion

Decreased cell stiffness is a prominent physical characteristic observed in malignant cells and particularly pronounced in tumor stem cells (1-6). Our results demonstrate that softer stiffness levels lead to diminished T cell responsiveness, as evident by smaller fractions of responding cells, reduced Ca^2+^ influx, impaired MTOC reorientation, and NFAT nuclear translocation. Our findings indicate that the inherent softening in tumor cells can significantly diminish corresponding T cell responses, substantially increasing their likelihood of evading immune surveillance and facilitating metastasis and the establishment of clones in distant locations.

How do T cells discern variations in target cell stiffness and respond to this mechanical cue? TCRs inherently respond to mechanical forces generated during engagement with antigens (41, 48), facilitating the exposure of more binding sites to high-affinity antigens (49, 50). In this context, when interacting with softer tumor cells, the forces generated from bound antigen-TCR complexes are likely smaller compared to those in interactions with stiffer tumor cells. This disparity could plausibly lead to decreased activation of TCR downstream signals including SOCE via Orai channels, thus influencing NFAT activation and its regulated genes to govern T cell fate. We show that inhibiting SOCE has no impact on the peak levels of [Ca^2+^]_int_ induced by CD3 antibody coating on substrates. However, when stimulated by soluble CD3 antibody, BTP-2-treated Jurkat T cells exhibit reduced [Ca^2+^]_int_ peak levels (45). These findings suggest that SOCE plays a substantial role in the initial Ca^2+^ influx triggered by global stimulation with soluble CD3 antibody, however, SOCE may not be a primary contributor to the initial Ca^2+^ influx induced by focal stimulation on substrates with physiological levels of stiffness.

To mount an effective immune response against tumor cells, T cells must swiftly engage with recognized target cells and form a functional IS to exert their effector function. MTOC reorientation towards the IS is crucial for T cells to exert their effector functions, as MTOC reorientation plays a pivotal role in both sustaining T cell signaling (14) and facilitating effective directional delivery of vesicles containing cytotoxic proteins or cytokines (51, 52). Regarding MTOC repositioning upon IS formation, our results show that Piezo1-regulated Ca^2+^ influx plays a key role, establishing a link between Piezo1-mediated mechanosensing and MTOC repositioning. It is reported that MTOC reorientation relies on dynein activity, particularly the formation of dynein ring-like structures at the IS (10, 53). MTOC translocation is significantly impaired when dyneins fail to form ring-like structures following IS formation (53). Although CRAC channel-regulated Ca^2+^ influx in T cells is proven to be essential for various key processes during IS formation including actin reorganization (54), our results show that blocking CRAC channels does not alter activation-triggered MTOC repositioning, suggesting that MTOC reorientation is independent of SOCE-regulated actin reorganization.

Besides rapid responses like MTOC reorientation, subsequent long-term responses are also essential for T cell function, including cytokine production, checkpoint molecule expression, and cell fate determination involving activation, proliferation, and differentiation (55, 56), which together define the potential of subsequent reactions and the destiny of T cells. These responses are regulated by activation and nuclear translocation of the corresponding transcription factors. NFAT proteins play a central role in T cell commitment. Our results show that for Jurkat T cells, CD3 antibody stimulation-induced translocation of NFAT into the nucleus is stiffness dependent and regulated by CRAC/Orai-mediated Ca^2+^ influx, but independent of Piezo1. A recent study has reported that fluid shear stress potentiates activation of Jurkat T cells induced by CD3/CD28 antibody-coated beads as evidenced by several early- and late-stage activation markers including NFAT nuclear translocation (38). One landmark of this shear stress-enhanced T cell activation, ZAP70 phosphorylation, is primarily regulated by Piezo1 and can be abolished by removal of extracellular Ca^2+^ (38). Considering our findings, shear stress-potentiated NFAT translocation should be independent of Piezo1 but regulated by CRAC/Orai1 channels. Interestingly, in microglial cells, considered the major immune cell type in the brain, activation of NFAT proteins is also independent of Piezo1 (57). In many other cell types, however, we noticed that Piezo1-mediated mechanosensing is linked to activation of the NFAT family, including macrophages (58), osteoblasts (59), erythroblasts (60), colon cancer stem-like cells (61), and cardiomyocytes (62). This suggests that different cell types adapt their mechanosensing pathways, with or without Piezo1 involvement, to govern mechanical cue-regulated NFAT nuclear translocation and its related gene regulation.

Rapid responses directly dictate the short-term functionality of T cells during contacts. Subsequently, T cells need to decide whether to sustain or halt their effector function depending on the surrounding microenvironment. For example, if the initially recognized target cell is surrounded by healthy cells, T cells should promptly cease their effector functions to avoid jeopardizing innocent bystander cells. On the other hand, if numerous other target cells are present, T cells should continue their effector functions for efficient elimination. Dysregulation in long-term responses can lead to inappropriate overactivation or suppression of T cell functions, resulting in autoimmune diseases or cancer. Consequently, decoupling rapid and long-term responses is crucial. Our results suggest that T cells employ stiffness as a regulatory cue to coordinate rapid and long-term responses via Piezo1 and Orai1.

In summary, our findings highlight the central role of target cell stiffness in regulating T cell functionality, including both rapid responses and long-term functions. The Ca^2+^ influx triggered by TCR activation emerges as a crucial factor for stiffness sensing. Differential regulation of stiffness-regulated rapid and long-term responses to stiffness is provided by Piezo1 and Orai1, which selectively control distinct mechano- and Ca^2+^-dependent processes. In addition, our results suggest that mechanosensing may act as a gatekeeper mechanism, setting a threshold below which T cell functions are completely inhibited. As soon as this threshold is exceeded, the T cells are sensitized so that they can react quickly and robustly. This delicate balance could ensure that T cells remain quiescent in non-diseased situations, preventing overreactions to harmless stimuli. Finally, our results provide direct evidence of how tumor cells exploit this stiffness-regulated mechanism to evade immune surveillance. Our data thus support the hypothesis that targeting the mechanosensitivity of T cells could potentially improve the efficacy of immunotherapy (63, 64).

## Materials and Methods

### Antibodies and reagents

All chemicals not specifically mentioned are from Sigma-Aldrich (highest grade). The following antibodies or reagents were used: Anti-γ-Tubulin antibody (Merck, T5192), Anti-Phospho-Zap-70 (Tyr319)/Syk (Tyr352) (Cell Signaling Technology, 27175), Anti-PIEZO1 (Thermo Fisher Scientific, PA5-72974), Anti-Orai1 (Sigma-Aldrich, O8264), Anti-NFAT1 (Cell Signaling Technology, 4389), Alexa Fluor Plus 488 Goat anti-Rabbit IgG (H+L) Secondary Antibody (Thermo Fisher Scientific, A32731), Alexa Fluor™ 647 Goat anti-Rabbit IgG (H+L) Secondary Antibody (Thermo Fisher Scientific), Alexa Fluor™ 568 Goat anti-Rabbit IgG (H+L) Secondary Antibody (Thermo Fisher Scientific, A-11036), Alexa Fluor^®^ 647 anti-human human CD3 Antibody (BioLegend, 300416), Alexa Fluor^®^ 488 anti-human human CD3 Antibody (BioLegend, 300415), Biotin anti-human CD3 Antibody (BioLegend, 317320), Biotin anti-human CD28 Antibody (BioLegend, 302904), Alexa Fluor^®^ 647 anti-Tubulin-α Antibody (BioLegend, 627908), Alexa Fluor™ 568 Phalloidin (ThermoFisher, A12380), Alexa Fluor™ 488 Phalloidin (ThermoFisher, A12379), Fura-2/AM (Thermo Fisher Scientific, F1221), SPY650-tubulin (Spiochrome, SC503), Hoechst 33342 (ThermoFisher Scientific, H3570), fibronectin (Sigma-Aldrich, F4759), Poly-L-ornithine (Sigma-Aldrich, P3655).

### Cell culture and transfection

Jurkat T cells (Clone E6-1, from ATCC), Raji (from DSMZ) and NALM-6 cells (kindly provided by Adolfo Cavalié, Saarland University) were cultured in RPMI 1640 (ThermoFisher Scientific) supplemented with 10% fetal calf serum (ThermoFisher Scientific) and 1% penicillin–streptomycin (ThermoFisher Scientific). Primary human CD4^+^ T cells were isolated from peripheral blood mononuclear cells obtained from healthy donors using Human CD4^+^ T Cell Isolation Kit (Miltenyi Biotec) according to the manual and were stimulated with Dynabeads™ Human T-Activator CD3/CD28 (ThermoFisher Scientific) with 30 U/ml of recombinant human IL-2 (Miltenyi Biotec). Unstimulated human primary T cells were cultured in AIMV (ThermoFisher Scientific) with 10% fetal calf serum and 1% penicillin–streptomycin. All cells were cultured at 37°C, 5% CO_2_ in a humidified incubator.

Regarding siRNA transfection, Jurkat T cells were transfected either with siRNA against Piezo1 (ON-TARGETplus SMARTpool human Piezo1 siRNA) or control siRNA (Horizon Discovery, previously Dharmacon) or with siRNA against Orai1 (a mixture of two modified siRNAs of human Orai1 (65), or control RNA with SE Cell Line 4D-Nucleofector™ X Kit (Lonza). Jurkat T cells were used for experiments 24 h (for Piezo) or 60 h (for Orai1) post transfection.

### Fluorescence-based Ca^2+^ imaging with Fura-2-AM

For Ca^2+^ imaging, as described previously (65), Jurkat T cells were loaded with 1 μM Fura-2-AM in RPMI 1640 with 10% FCS at room temperature for 25 min. Then cells were washed with centrifugation, resuspended in 1 mM Ca^2+^ Ringer’s solution, and seeded on poly(acrylamide) (PAAm) hydrogel substrate if not otherwise mentioned. Afterwards, Ca^2+^ imaging is acquired immediately. Fluorescence was emitted by 340 nm or 380 nm and infrared images were taken every 5 sec for 25 min at room temperature. The captured images were analyzed by T.I.L.L. Vision software. The traces were analyzed with the software Igor Pro6. For the experiment with BTP-2, all solutions contain 10 μM BTP-2 or the vehicle. The time 0 is defined as the time when cells dock on the substrate. The cells showing elevated Ca^2+^ levels ([Ca^2+^]_int_ exceeding the threshold Ratio 340/380=0.3) were defined as responsive cells. The max Ca^2+^ increase rate is the maximum Ca^2+^ increase rate between 2 time points of measurement. The peak was defined as the maximum value of ratio 340nm/380nm. The plateau levels the average of 340nm/380nm ratios from the last 30 sec.

### Conjugation of T cells with target cells

Target cells (NALM-6 or Raji cells) were pulsed with staphylococcal enterotoxin A (SEA, 0.1 µg/ml) and SEB (0.1 µg/ml) at 37°C for 40 min. For conjugation, Jurkat T cells and pulsed target cells were mixed and settled on the fibronectin (0.1 mg/ml)-coated coverslip at 37°C for 15 min. To reduce the stiffness of target cells, following SEA/SEB pulsing, NALM-6 cells were loaded with EGTA-AM (500 µM) at 37°C for 30 min and then washed twice to remove unloaded EGTA-AM.

### Preparation and biofunctionalization of substrates

For Ca^2+^ imaging, poly (acrylamide-co-acrylic acid) (PAAm-co-AA) hydrogels with stiffness levels of 2, 12, and 50 kPa were prepared and functionalized following a previously described protocol (41). Briefly, a mixture of AAm monomer and bis-AAm crosslinker was prepared with varying ratios, while maintaining a constant ratio of AA. Hydrogel discs were formed between two glass coverslips. The PAAm-co-AA hydrogels were firstly functionalized with biotin-PEG8-NH2 through an EDC/NHS activation step as follows. The PAAm-co-AA film was covered with 100 µL EDC/NHS solution (39 mg/12 mg in 0.1 M, pH 4.5 MES buffer) for 15 min, washed thoroughly with PBS and then directly incubated with 100 µL of biotin-PEG8-NH2 (1 mg/mL) solution in a petri dish for 2 hr at RT. The functionalized hydrogels underwent a triple wash with PBS and were stored in PBS at 4°C until use. Subsequently, the hydrogels of varying stiffness were incubated with streptavidin solution (100 µL, 100 μg/mL) for 1 –1.5 h (2 kPa) or 2.5 –3 h (12 and 50 kPa). These hydrogels were then subjected to overnight incubation with biotinylated anti-CD3 antibody (100 μg/mL, 30 μL) or biotinylated anti-CD28 antibody (100 μg/mL, 30 μL) at 4°C.

For immunostaining and live-cell imaging, CytoSoft® Imaging 96-Well plates with stiffness of 2, 16 and 64 kPa from Advanced BioMatrix were used if not otherwise mentioned. Plates were coated with biotinylated anti-CD3 antibody or anti-CD28 antibody (30 μg/mL, 50 μL/well) in PBS at 4°C for overnight. Afterwards, the antibody solution was removed and washed with PBS 3 times. For examining coating efficiency, Alexa Fluor^®^ 488-conjugated anti-CD3 antibody (30 μg/mL, 50 μL/well) were used to coating the wells. The antibody density on the substrate was assessed the fluorescence intensity of Alexa Fluor^®^ 488 with CellDiscover7 (20× objective, NA:0.7).

### Fluorescence-based imaging with MTOC translocation

Jurkat T cells were loaded with SPY650-tubulin (1:2000) in RPMI 1640 with 10% FCS at 37°C, 5% CO_2_ in a humidified incubator for 25 min. 1 μM GsMTx-4 or the vehicle is present in all solutions. After staining, cells were washed with centrifugation. Cells were mixed with 1 mM Ca^2+^ Ringer’s solution and plated into biotin anti-human CD3 Antibody coated CytoSoft® Imaging 96-Well Plate. Afterwards, images were acquired with a Zeiss Cell Observer HS system with a 20× alpha objective and an AxioCam MRm Rev. 3. During the experiment, the fluorescence of tubulin was emitted by 625 nm and acquired with a Cy5 (F36-523) filter set with 1 μm z-step size. The images were acquired every 9 sec at room temperature. The position of MTOC was defined with TrackMate of Fiji Plugins.

### Epifluorescence deconvolution microscopy and immunofluorescence

For substrates, T cells were settled on antibody-coated substrates at 37°C for the indicated periods of time. For conjugation, T cells were settled with SEA/SEB-pulsed target cells (NALM-6 cells or Raji cells) on fibronectin (0.1 mg/ml)-coated coverslips at 37°C for 15 min. Cells were then fixed with 4% paraformaldehyde and permeabilized with 0.3% Triton-100 except for γ-tubulin antibody (permeabilized with methanol/acetone solution at -20°C for 20 min. After immune staining, the fixed samples were measured with a Zeiss Cell Observer HS system containing GFP (38HE; Zeiss), ET-Texas Red (Chroma Technology), DAPI HE (Zeiss), Texas Red HE (Zeiss), and Cy 5 HE (Zeiss) filter sets. The samples on PAAm hydrogel were measured with alpha Plan-Apochromat 63x/1.40 Oil DIC M27. The samples on the coverslip were measured with alpha Plan-Apochromat 63x/1.40 Oil DIC M27. The samples in CytoSoft® Imaging 96-Well Plates were measured with Fluar 40x/1.30 Oil M27. The acquired images were preceded for deconvolution with the Huygens Essential program by using a theoretical point spread function.

### Immunostaining and flow cytometry

Samples were fixed with 4% paraformaldehyde (PFA) and then washed twice with PBS/0.5% BSA. Cells were permeabilized and blocked with 0.3% Triton-100 in PBS containing 5% FCS and 0.5% BSA for 30 min at room temperature. And then cells were stained with the indicated primary antibody for 1 hr at room temperature followed by staining of Alexa Fluor™ 647 labeled secondary antibody. Flow cytometry data were acquired using a FACSVerse™ flow cytometer (BD Biosciences) and were analyzed with FlowJo v10 (FLOWJO, LLC).

### Real-time deformability cytometry (RT-DC)

To assess the stiffness of NALM-6 cells RT-DC (Zellmechanik Dresden) was used (40). NALM-6 cells were either treated with DMSO or EGTA-AM for 30 min after which they were resuspended in 100 µl of Cell Carrier B solution (phosphate-buffered saline with the addition of long-chain methylcellulose polymers of 0.6 w/v%). A microfluidic PDMS chip with a 300 µm long central constriction of 30 µm×30 µm cross-section was assembled on the stage of an inverted microscope (Zeiss). The cell suspension was loaded on the chip using a syringe pump. The cells flowing through the microfluidic channel deform due to the shear stresses and pressure gradient caused by the flow profile (40). Each event is imaged live using a CMOS camera and analyzed in real-time. Events were acquired at a flowrate of 0.12 µl/s. The mechanical properties of the cells were analyzed using ShapeOut 2 (Zellmechanik Dresden) which employs linear mixed models to calculate statistical significance.

### Statistical Analysis

Data are presented as mean. GraphPad Prism 6 Software (San Diego, CA, USA) was used for statistical analysis. If the number of data points is smaller than 8, the differences between two columns were analyzed by the student’s t-test with two tails. Otherwise, the data were first examined for Gaussian distribution. If the dataset fits Gaussian distribution, the differences between two columns were analyzed with the student’s t-test with two tails, otherwise with the Mann-Whitney test with two tails. the differences between three columns were analyzed by the One-way ANOVA or nonparametric test with two tails. If datasets fit Gaussian distribution, the differences between three columns were analyzed with a one-way ANOVA with Tukey’s multiple comparisons, otherwise the differences between three columns were analyzed with the Krushal-Wallis test with Dunn’s multiple comparisons. The RT-DC data were analyzed with linear mixed-effect models.

## Ethical Considerations

Research carried out for this study with healthy donor material (leukocyte reduction system chambers from human blood donors) is authorized by the local ethic committee [declaration from 16.4.2015 (84/15; Prof. Dr. Rettig-Stürmer)].

## Acknowledgments

We thank Prof. Hermann Eichler (Institute of Clinical Hemostaseology and Transfusion Medicine) for blood samples; Carmen Hässig, Cora Hoxha, Gertrud Schäfer, Sandra Janku, and Kathleen Seelert for excellent technical help; Britta Abt and Jennifer Kasper for helping in hydrogels; Adolfo Cavalié for providing NALM-6 cells; Dalia Alansary for Ca^2+^ imaging, Orai1 siRNA transfection; Galia Montalvo and Franziska Lautenschläger for their kind assistance in RT-DC; Hong Zhang for helping in immunostaining. This project was funded by the Deutsche Forschungsgemeinschaft, SFB 1027 (Projects A2 (BQ), A11 (MH), and B6 (AdC) and “Large Equipment Grants” (GZ: INST 256/419-1 FUGG for the lightsheet microscope, GZ: INST 256/423-1 FUGG for the flow cytometer, GZ: INST 256/429-1 FUGB for ImageXpress, and GZ: INST 256/555-1 FUGG for the Celldiscoverer 7), mini-proposal of SFB 1027 (to R.Z.).

