## Supplementary Information for "T cell polarization and NFAT translocation are stiffness-dependent and are differentially regulated by Piezo1 and Orai1"

### Supplementary Figures

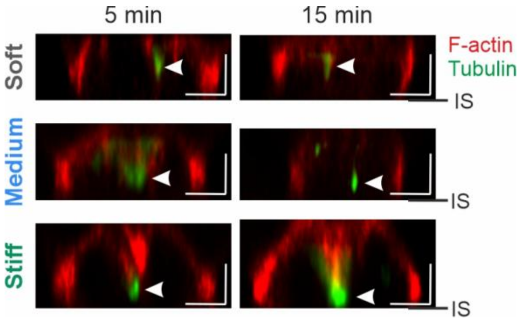


**Figure S1. Stiffness influences MTOC reorientation in Jurkat T cells.** Jurkat T cells were settled on functionalized substrates for varying time periods at 37°C before fixation. MTOC was stained with the anti-Tubulin antibody and the position of the IS was identified with Phalloidin-labeled actin. MTOC was indicated with the white arrow. 96-well CytoSoft plates with stiff levels of 2 kPa (soft), 16 kPa (medium), and 64 kPa (stiff) were used. Scale bars are 5 μm.

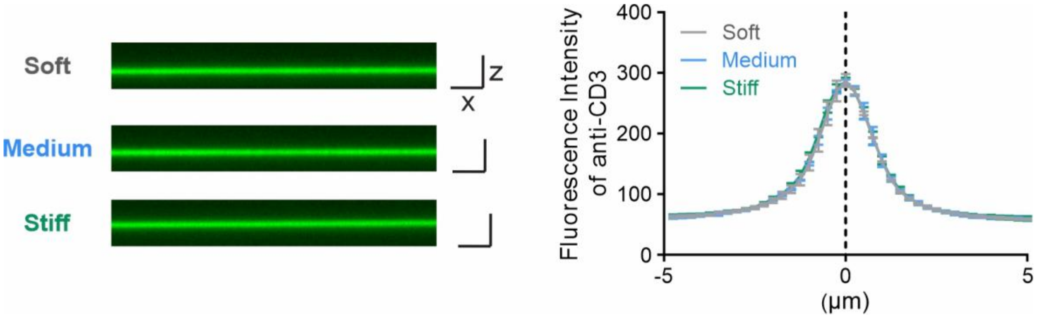


**Figure S2. The coating efficiency is comparable for all three stiffness levels**. Alexa Fluor 488 anti-human CD3 Antibody (30 μg/ml in PBS) was coated on bottoms of wells in CytoSoft plates with stiffness levels of 2 kPa (soft), 16 kPa (medium), and 64 kPa (stiff). The fluorescence of the antibody was measured with 20× objective (NA: 0.7) using the confocal mode of CellDiscover 7. The step size of the z stack is 250 nm. Quantification (right panel) is from 3 independent experiments and is shown as the mean ± SEM. Scale bars are 10 μm.

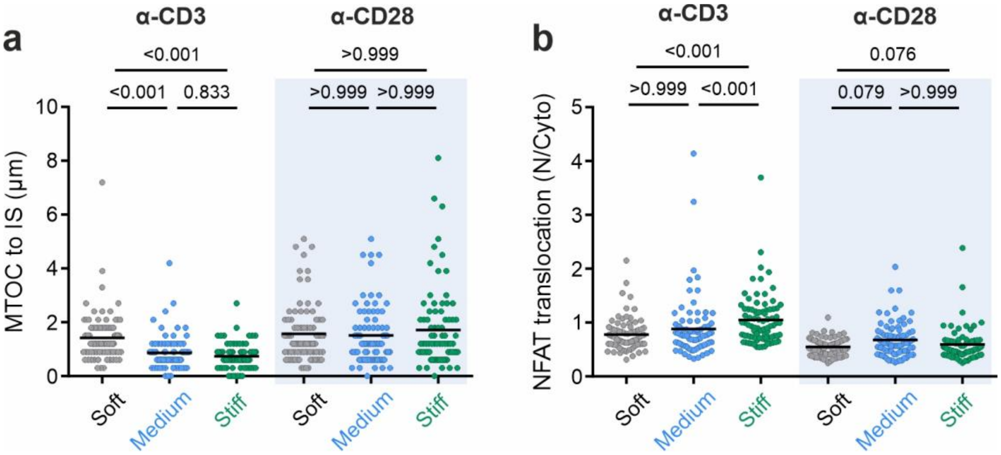


**Figure S3. Substrates functionalized CD28 antibody did not induce MTOC reorientation or NFAT nuclear translocation**. CD28 antibody-functionalized silicon surface cannot induce stiffness-dependent MTOC reorientation (**a**) and NFAT translocation (**b**). T cells were plated on the antibody-functionalized surface at 37°C for 15 min before fixation. MTOC was stained with the anti-tubulin antibody and the position of the IS was identified with Phalloidin-labeled actin (**a**). NFAT and the nucleus were stained with anti-NFAT antibody and Hoechst 33342, respectively (**b**). A one-way ANOVA with Tukey's multiple comparisons was conducted for **a**, **b.** 96-well CytoSoft plates with stiff levels of 2 kPa (soft), 16 kPa (medium), and 64 kPa (stiff) were used.

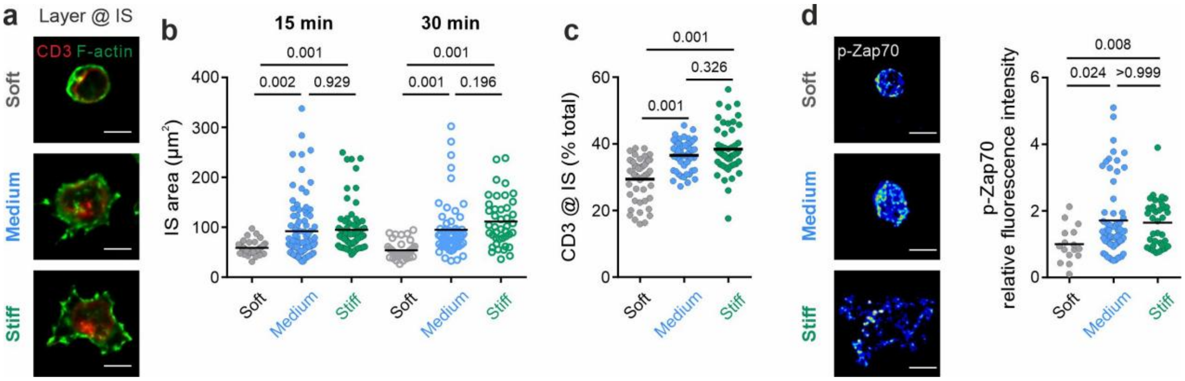


**Figure S4. T cells are activated by functionalized hydrogels.** Jurkat T cells were settled on CD3 antibody-functionalized hydrogels with 2 kPa (soft), 12 kPa (medium), and 50 kPa (stiff) at 37°C for 15 min or 30 min before fixation. (**a-b**) T cells spread and CD3 accumulates upon contact. F-actin was used to identify the IS position and the contact area. Exemplary cells at 30 min are shown in **a**. Quantification of individual cells at 15 and 30 min is shown in b. Distribution of CD3 is quantified at 30 min (**c**). The proximity to the IS region was defined as the region from the contact interface up to one-quarter of cell height. (**d**) Jurkat T cells were fixed after 15 min. Phosphorylation of Zap70 is enhanced upon contact. Exemplary cells with pseudo color of phosphorylated Zap70 (p-Zap70) staining. Quantification of individual cells is shown in the right panel. Scale bars are 5 μm. A one-way ANOVA with Tukey's multiple comparisons was conducted for **b** and **c**. A Krushal-Wallis test with Dunn's multiple comparisons was conducted for **d**. Results are from two (**b**) or three (**a**, **c**, **d**) independent experiments.

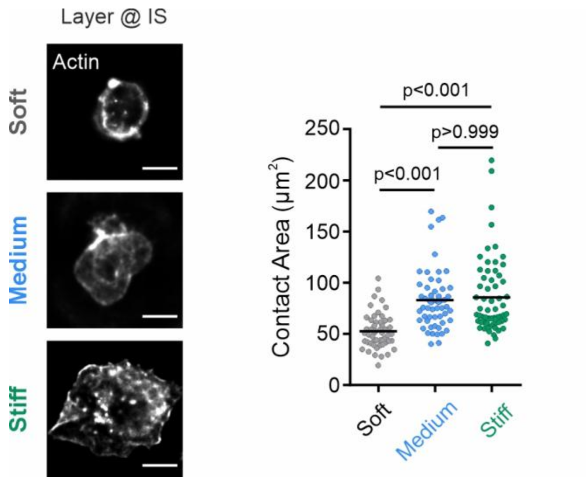


**Figure S5. Anti-CD28 antibody functionalized hydrogels induce contact area of T cells.** Jurkat T cells were settled on CD28 antibody-functionalized with stiffness levels of 2 kPa (soft), 12 kPa (medium), and 50 kPa (stiff). T cells spread upon contact. F-actin was used to identify the IS position and the contact area. T cells were plated on anti-CD28 antibody-functionalized surface at 37°C for 15 min before fixation. A Krushal-Wallis test with Dunn's multiple comparisons was conducted.

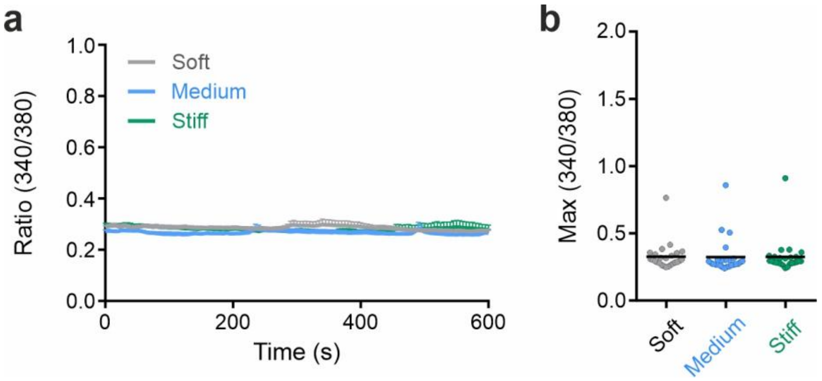


**Figure S6. Anti-CD28 antibody functionalized hydrogels do not induce $Ca^{2+}$ influx in T cells.** Anti-CD28 antibody-functionalized hydrogels do not trigger a $Ca^{2+}$ rise in T cells. Jurkat T cells were loaded with Fura-2-AM and live cell images were acquired every 5 sec for 25 min at RT. The $[Ca^{2+}]_{int}$ in T cells is shown after the cell settles down on the surface. Mean $[Ca^{2+}]_{int}$ values of cells are shown in **a**, in which time 0 is defined as the time point when T cells settle down on the surface. $Ca^{2+}$ max ratio (340/380) in 600 sec (**b**) was shown in the figure. Results are shown as mean ± SEM in (**a**) or as mean (**b**).

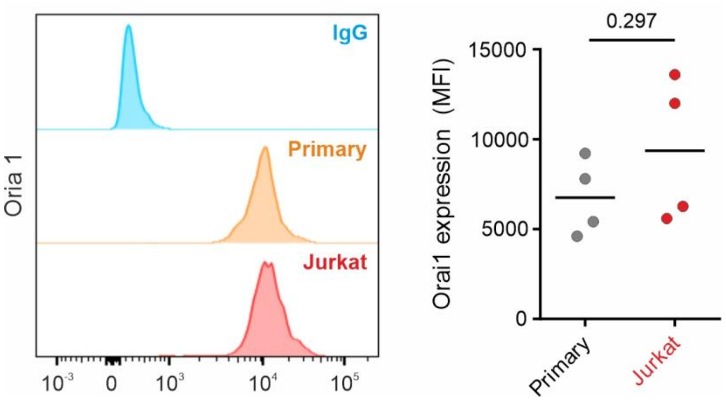


**Figure S7. Expression of Piezo1 is similar in stimulated human primary CD4⁺ T cells and Jurkat T cells.** Expression of Piezo1 in T-Activator CD3/CD28 antibody beads stimulated human primary $CD4^+$ T cells and Jurkat T cells were measured with flow cytometry. Results are from four independent experiments.

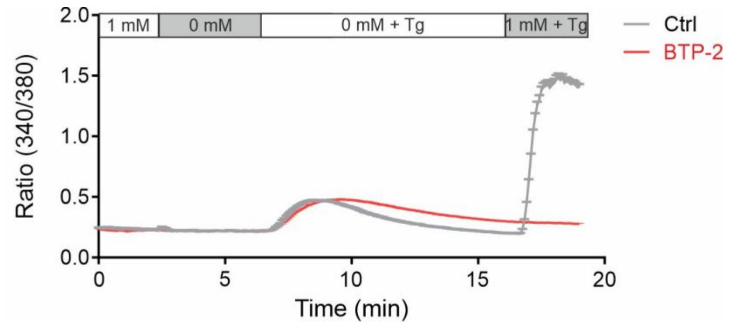


**Figure S8. BTP-2 treatment abolished the function of CRAC channels**. Jurkat T cells loaded with Fura-2-AM were settled on poly-L-ornithine-coated coverslips in 1 mM $Ca^{2+}$ ringer solution. Then ringer solutions with or without thapsigargin (TG) were perfused as shown in the stretch. DMSO or BTP-2 (10 μM) was present in all solutions of the experiment. DMSO: 196 cells, BTP-2: 233 cells. Data are shown in mean ± SEM from two independent experiments.

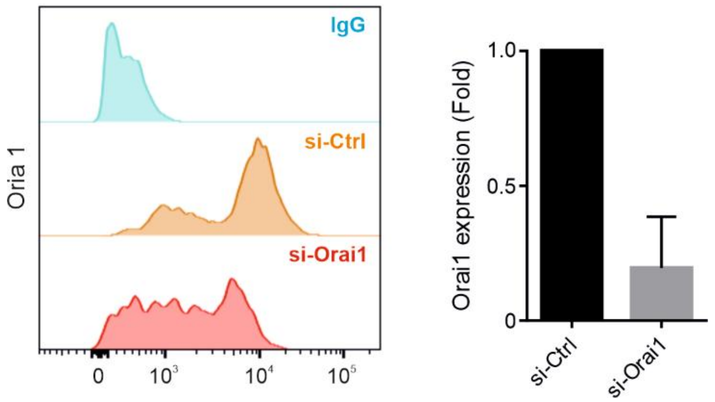


**Figure S9. Down-regulation of Orai1 by siRNA**. Jurkat T cells were transfected with control siRNA (si-Ctrl) or siRNA targeting Orai1 (si-Orai1). Transfected cells were fixed 60 hours after transfection, and were stained with Orai1 antibody. The expression of Orai1 was measured with flow cytometry. One representative experiment is shown in the left panel. Quantification of fold change in mean fluorescence intensity is shown in the right panel. The results are from two independent experiments.

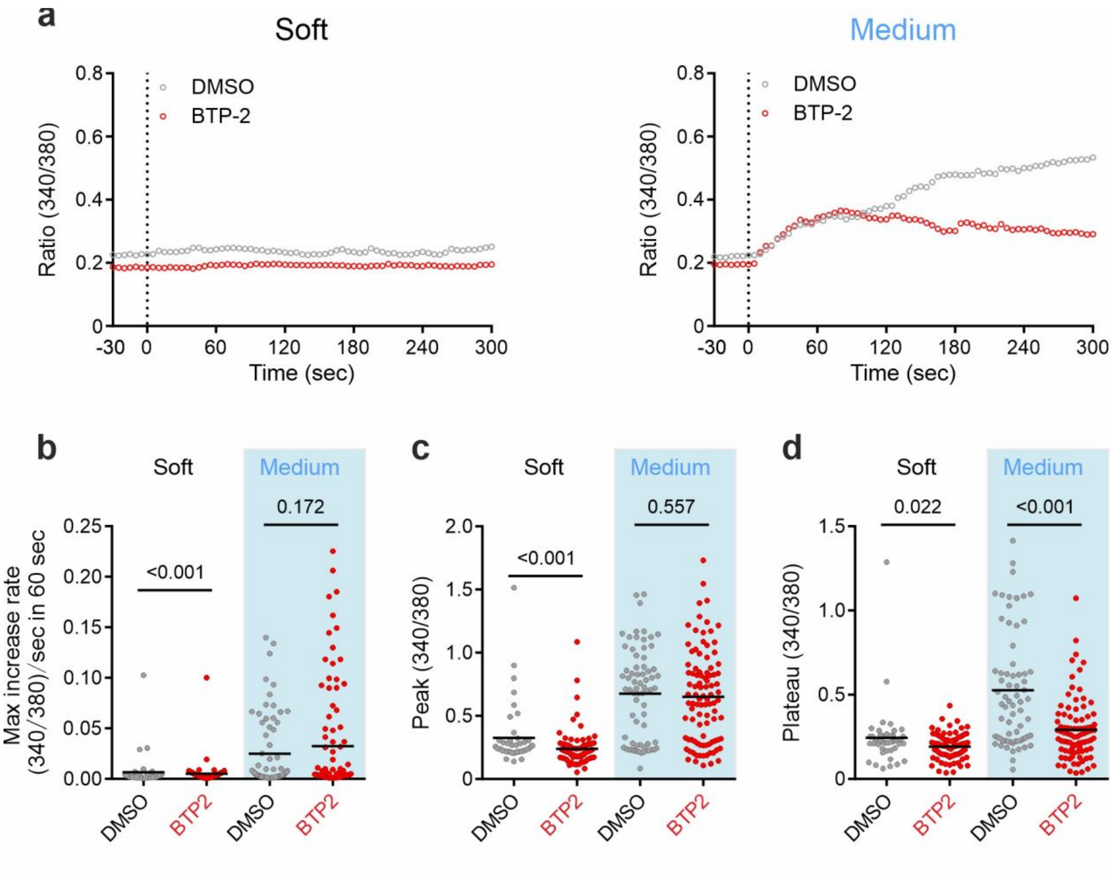


**Figure S10. Impact of Orai1 channels on TCR activation-triggered $Ca^{2+}$ rise on the soft and medium surface**. T cells were settled on different stiffness substrates (soft: 2 kPa hydrogel, medium: 12 kPa hydrogel). The mean values of all cells for each condition are shown in **a**, in which time 0 is defined as the time point when T cells settle down on the surface. $Ca^{2+}$ max increase rates (**b**), peak (**c**), and plateau (**d**) were quantified. The results of DMSO treatment on 2 kPa hydrogel are from 2 independent experiments. Other results are from at least three independent experiments. A Mann-Whitney test was conducted for **b**-**d**.

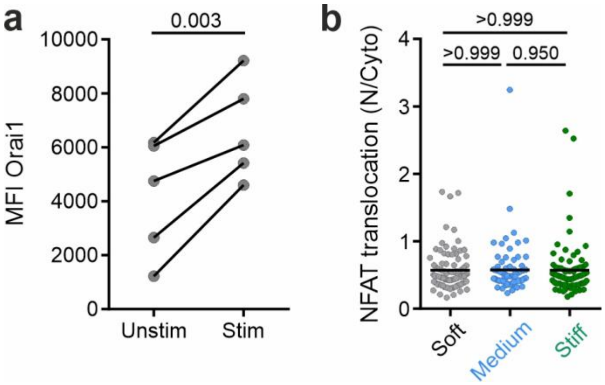


**Figure S11. NFAT nuclear translocation does not depend on stiffness in unstimulated human primary CD4⁺ T cells.** (**a**) ORAI1 expression lever is lower in unstimulated human primary $CD4^+$ T cells than T-Activator CD3/CD28 antibody beads stimulated human primary $CD4^+$ T cells from the same donor. The expression of Orai1 was measured with flow cytometry. Data are from 5 donors and analyzed with the paired Student's t-test. (**b**) In unstimulated human primary CD4+ T cells, NFAT nuclear translocation does not depend on stiffness. T cells were plated on anti-CD3 antibody-functionalized surface at 37°C for 15 min before fixation. NFAT and the nucleus were stained with anti-NFAT antibody and Hoechst 33342, respectively. 96-well CytoSoft plates with stiff levels of 2 kPa (soft), 16 kPa (medium), and 64 kPa (stiff) were used. Data are from 3 donors and analyzed with the Krushal-Wallis test with Dunn's multiple comparisons.

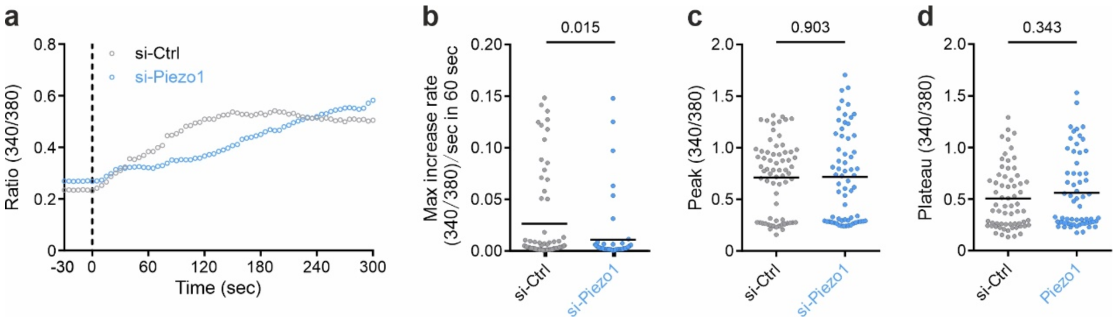


**Figure S12. Piezo1 regulates $Ca^{2+}$ increase rates on medium stiffness substrates.** Impact of Piezo1 on TCR activation-triggered $Ca^{2+}$ rise. Jurkat T cells transfected with siRNA were loaded with Fura-2-AM and then settled on anti-CD3 antibody functionalized medium stiffness of hydrogels (12 kPa). Averaged curves of cells for each condition are shown in **a**, in which time 0 is defined as the time point when T cells settle down on the surface. $Ca^{2+}$ max increase rate (**b**), peak (**c**), and plateau (**d**) were quantified for individual cells. A Mann-Whitney test was conducted for **d-f**. All results are from three independent experiments. 96-well CytoSoft plates with stiffness levels of 2 kPa (soft), 16 kPa (medium), and 64 kPa (stiff) were used.

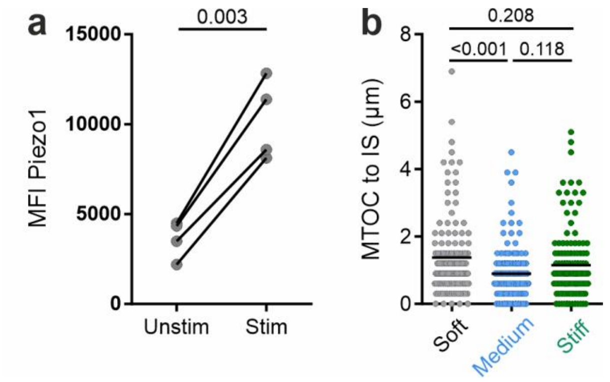


**Figure S13. MTOC reorientation does not depend on stiffness in unstimulated human primary CD4+ T cells.** (**a**) Piezo1 expression lever is lower in unstimulated human primary CD4+ T cells than T-Activator CD3/CD28 antibody beads stimulated human primary CD4+ T cells from the same donor. The expression of Orai1 was measured with flow cytometry. Data are from 4 donors and analyzed with the paired Student's t-test. (**b**) In unstimulated human primary CD4+ T cells, MTOC reorientation does not depend on stiffness. T cells were plated on anti-CD3 antibody-functionalized surface at 37°C for 15 min before fixation. MTOC was stained with the anti-tubulin antibody and the position of the IS was identified with Phalloidin-labeled actin. 96-well CytoSoft plates with stiff levels of 2 kPa (soft), 16 kPa (medium), and 64 kPa (stiff) were used. Data are from 4 donors and analyzed with the Krushal-Wallis test with Dunn's multiple comparisons.
